# Role of Dopamine Neurons in Familiarity

**DOI:** 10.1101/2023.10.25.564006

**Authors:** Sixtine Fleury, Rhonda Kolaric, Justin Espera, Quan Ha, Jacquelyn Tomaio, Ulrik Gether, Andreas Toft Sørensen, Susana Mingote

## Abstract

Dopamine neurons signal the salience of environmental stimuli, influencing learning and motivation. However, research has not yet identified whether dopamine neurons also modulate the salience of memory content. Dopamine neuron activity in the ventral tegmental area (VTA) increases in response to novel objects and diminishes as objects become familiar through repeated presentations. We proposed that the declined rate of dopamine neuron activity during familiarization affects the salience of a familiar object’s memory. This, in turn, influences the degree to which an animal distinguishes between familiar and novel objects in a subsequent novel object recognition (NOR) test. As such, a single familiarization session may not sufficiently reduce dopamine activity, allowing the memory of a familiar object to maintain its salience and potentially attenuating NOR. In contrast, multiple familiarization sessions could lead to more pronounced dopamine activity suppression, strengthening NOR. Our data in mice reveals that, compared to a single session, multiple sessions result in decreased VTA dopamine neuron activation, as indicated by c-Fos measurements, and enhanced novelty discrimination. Critically, when VTA dopamine neurons are chemogenetically inhibited during a single familiarization session, NOR improves, mirroring the effects of multiple familiarization sessions. In summary, our findings highlight the pivotal function of dopamine neurons in familiarity and suggest a role in modulating the salience of memory content.

## Introduction

As organisms navigate complex and dynamic environments, it is essential to discriminate between both familiar and novel events. Familiarity sets the boundaries of an animal’s existing knowledge, facilitating the detection of new events (Brown and Aggleton, 2001). Novelty triggers alerting responses to unexpected information, aids in initiating new learning, and enhances the formation of new memories (Tulving and Kroll, 1995; Bromberg-Martin et al., 2010; Duszkiewicz et al., 2019; Morrens et al., 2020). Brain circuits controlling novelty and familiarity detection are subjects of intensive research not only for their importance in adaptive behavior, but also due to their decline in aging and Alzheimer’s disease (Brown and Xiang, 1998; Eichenbaum et al., 2007; Kafkas and Montaldi, 2014, 2018; Bastin et al., 2019; Chen et al., 2020; Tapper and Molas, 2020).

Research across drosophila, rodents, and humans consistently points to the involvement of dopamine neurons in modulating responses to novelty and the shift toward familiarity (Horvitz, 2000; Bunzeck and Düzel, 2006; Bromberg-Martin et al., 2010; Hattori et al., 2017; Morrens et al., 2020). Rodents innately explore novel objects within their surroundings, and their exploratory behavior wanes upon repeated exposure to the same object. As this shift from novelty to familiarity occurs, dopamine neurons in the ventral tegmental area (VTA) show increased activity in response to novel objects and reduced activation during subsequent encounters (Gunaydin et al., 2014). Interestingly, while novelty triggers increased exploration and dopamine neuron activity, the chemogenetic or optogenetic inhibition of VTA dopamine neurons during the introduction of a new object does not disrupt exploratory behavior (Gunaydin et al., 2014; Bariselli et al., 2018). These findings indicate that dopamine neuron activity does not control the ongoing exploratory behavior but might influence future learning and memory.

There is strong evidence that dopaminergic responses influence learning and motivation by signaling the perceived salience of stimuli in the environment (Berridge, 2007; Flagel et al., 2011; Morrens et al., 2020; Kutlu et al., 2021, 2022). In associative learning tasks, animals form associations between cues and outcomes and develop conditioned responses to these cues. These conditioned responses are acquired more slowly for familiar, non-salient cues associated with lower dopamine neuron activation than for novel cues that elicit higher dopamine neuron activity. Slower learning for familiar cues is known as the “latent inhibition effect.” In this learning paradigm, dopamine influences the perceived salience of the cue. Optogenetically inhibiting dopamine neurons during the presentation of a new cue reduces its salience and induces the latent inhibition effect (Morrens et al., 2020; Kutlu et al., 2022). It remains unclear whether dopamine neurons influence the memory of previously encountered objects in the same manner. This function would help animals form memories as either salient and familiar or familiar but non-salient.

Memory of previously encountered objects in mice is frequently assessed using the novel object recognition (NOR) paradigm (Ennaceur, 2010). This paradigm consists of a sample phase and a test phase. During the sample phase, animals undergo familiarization sessions where they are repeatedly presented with two identical objects. In the test phase, the animals are subjected to a novelty object recognition (NOR) session in which one of the familiar objects is replaced with a new one. An increase in time spent exploring a new object, compared to a familiar one, is considered indicative of the ability to remember previously encountered objects and to form what is known as recognition memory (Brown and Banks, 2015). However, successful discrimination between novel and familiar objects is also influenced by factors such as the perceived salience of previously encountered objects. In a recent novelty preference study using male mice, adding a female odor enhanced an object’s salience (Hornoiu et al., 2020). When the animals were introduced to this highly salient but familiar object alongside a new odorless object, they explored both nearly equally, indicating no preference between the novel object and the familiar but salient object.

We hypothesized that the activity of dopamine neurons during the familiarization phase of the NOR dictates the salience linked to the memory of the familiar object. This function in familiarization subsequently impacts performance in the following novelty discrimination test. In formulating our hypothesis, we relied on the well-established observation that repeated presentations of the same object gradually diminish the object’s salience and novelty-induced dopaminergic responses (Gunaydin et al., 2014; Bariselli et al., 2018). A brief, single familiarization session might lead to minor reductions in dopamine neuron activity and be insufficient to diminish the salience of the familiar object. Thus, in subsequent novelty discrimination tests, the familiar object would be perceived as equally salient as the novel one, leading to a weaker NOR. In contrast, we expect that multiple familiarization sessions result in more pronounced reductions in both dopamine neuron activity and the salience of the familiar stimulus, culminating in a successful NOR. To evaluate the role of dopamine neurons in familiarity and its causal influence in determining the salience of memories of familiar objects, we posed three questions: (i) Does a single familiarization session produce weaker discrimination compared to multiple sessions? (ii) Are dopamine neurons more active during the first familiarization session than after multiple sessions? (iii) If we inhibit dopamine neurons during a single familiarization session, will it enhance novelty recognition, mirroring the effects of multiple familiarization sessions? Our results suggest that dopamine neurons significantly influence familiarity by modulating the salience of memory content.

## Materials and Methods

### Subjects

Mice were maintained following the guidelines of the NIH Guide for the Care and Use of Laboratory Animals under protocols approved by the CUNY ASRC Comparative Medicine Unit (CMU). Mice were co-housed and acclimatized with ad libitum food, water, and enrichment materials. Mice were exposed to a 12 h light/dark cycle, and experiments were conducted during the light cycle and after three handling sessions (10 min). The first experiment used C57BL6J male mice aged 3-5 months, and subsequent experiments involved male heterozygous tyrosine hydroxylase (TH)-2A-Flpo mice aged 4-6 months (MMRRC Repository, #050618-MU). All mice were bred and raised in our CMU.

### Stereotaxic Surgery and AAV Injection

Anesthetized TH-2A-Flpo mice (isoflurane 2%) underwent bilateral injections of AAV8-hSyn-fDIO-hM4Di-mCherry (4×1012 titer) into the VTA (0.7μl; coordinates relative to bregma: AP −3.3 mm, DV −4.3 mm, LM −0.5 mm). Mice in the mock CNO (Clozapine-N-Oxide) group received intracranial injections of sterile phosphate-buffered saline (PBS). The Flp-dependent inhibitory Designer Receptors Exclusively Activated by Designer Drugs (DREADD) virus was made by collaborator Andreas Toft Sørensen and validated in Ulrik Gether’s laboratory (Runegaard et al., 2018). Injections were performed using Nanoject II (Drummond Scientific). Animals were allowed to recover for four weeks before behavioral testing.

### Drug preparation and injection

CNO (Millipore Sigma, Cat# SML2304) was diluted in saline (0.9% sodium chloride) and injected intraperitoneal (i.p.) at a volume of 10ml/kg.

### Novel Object Recognition (NOR) Procedure

Mice were habituated to open field arenas (40×40×40cm; MazeEngineers, USA) for 30 minutes on the first day and 15 minutes on the second day. In the single familiarization session protocol, animals were exposed to one 20-minute familiarization session and allowed to explore two identical and north-placed objects (soda cans). Mice were individually housed for 30 minutes before undergoing a 10-minute NOR test, during which one object was replaced with a new one (a salad dressing bottle). The multiple familiarization sessions protocol exposed mice to three familiarization sessions across three days. In the chemogenetic experiment, mice got two rounds of the single familiarization session protocol, 14 days apart, with counterbalanced treatment order (see schematic in **Figure 3B**). Half received i.p. injections of CNO (1mg/kg) or saline 30 minutes before familiarization in the first round and before NOR in the second round. The other half had the reverse order of treatment. To prevent habituation from first-round experiences, we covered the arena floors with orange plastic sheet in the second round; objects were positioned to the east, and the familiar objects were round, transparent water bottles, while the novel object was a square, dark iced tea bottle.

### Behavioral Data Analysis

A video camera was positioned above the open field, and video tracking was performed using EthoVision XT 17 (Noldus, USA). We analyzed the distance traveled and time exploring objects using GraphPad Prism 9. Object exploration was quantified when the mouse’s nose was within a 2cm radius around the object. We calculated a discrimination index for exploration time using the formula: (Novel object – Familiar object) / (Novel object + Familiar object).

### Immunohistochemistry (IHC) and Imaging

Post-anesthesia (Ketamine 90 mg/kg + Xylazine 10mg/kg) mice underwent transcardial perfusion with cold PBS followed by 4% paraformaldehyde. Extracted brains were sectioned using a vibrating microtome (DSK MicroSlicer ZERO 1N) and stored in cryoprotectant. For IHC, sections underwent three PBS washes, incubated in glycine (100 mM) for 30 minutes, rewashed, blocked with 10% normal goat serum (Millipore) in 0.1% PBS-T solution for 2 hours, and were incubated with primary antibodies for 48 hours at 4°C. Primary antibodies included anti-mCherry (1:5000 dilution, rat monoclonal, Invitrogen #M11217), anti-c-Fos (1:5000, rabbit polyclonal, Abcam #190289), and anti-TH (1:2000, chicken polyclonal, Aves Labs, TYH-0020). After PBS washes, sections were incubated in secondary antibodies for 45 minutes at room temperature, which included anti-rat Alexa Fluor 568 (1:200 dilution, Invitrogen Cat# A1107), anti-rabbit Alexa Fluor 647 (1:200 dilution, Invitrogen Cat# A32733), and anti-chicken Alexa Fluor 488 (1:200 dilution, Invitrogen Cat# A32931). After a final wash, sections were mounted in slides using Prolong Gold (Invitrogen) and imaged with a Leica DM6 B epifluorescence microscope. We used the ImageJ cell counter plugin to count labeled neurons.

### Statistical analysis

Data were analyzed using two-tailed paired and unpaired t-tests, one-way-ANOVA, or two-way repeated measures ANOVA, as indicated in the results section, using Graph Prism 9. If the data set did not pass the normality test (Shapiro-Wilk test), we used non-parametric statistics (Wilcoxon matched-pairs signed rank test and Kruskal-Wallis test). Statistical significance was set at P < 0.05, and data in graphs were presented as averages and standard error of the mean. Estimation plots were used to show the data analyzed using t-tests and help visualize the magnitude of the effect size and precision. The effect size was represented as the difference between means, and the precision of the calculated effect size was expressed as 95% confidence intervals.

## Results

### A single familiarization session produces weaker novelty discrimination than multiple familiarization sessions

We examined the impact of the number of familiarization sessions on NOR using male C56BL6J mice. Mice were divided into two groups: the first group experienced a single familiarization session (n=11, **Figure 1A**), while the second group was exposed to 3 familiarization sessions (n=10, **Figure 1B**) before the NOR test. Over these multiple sessions, there was a consistent decline in time spent exploring identical objects (**Figure 1C**). A repeated measures two-way ANOVA revealed a significant main effect of the number of familiarization sessions (F_(2, 36)_ = 3.329; *P* = 0.047) with no significant effects of object location or significant interaction. These results suggest that animals become habituated to repeated presentations of the same objects.

**Figure 1.**
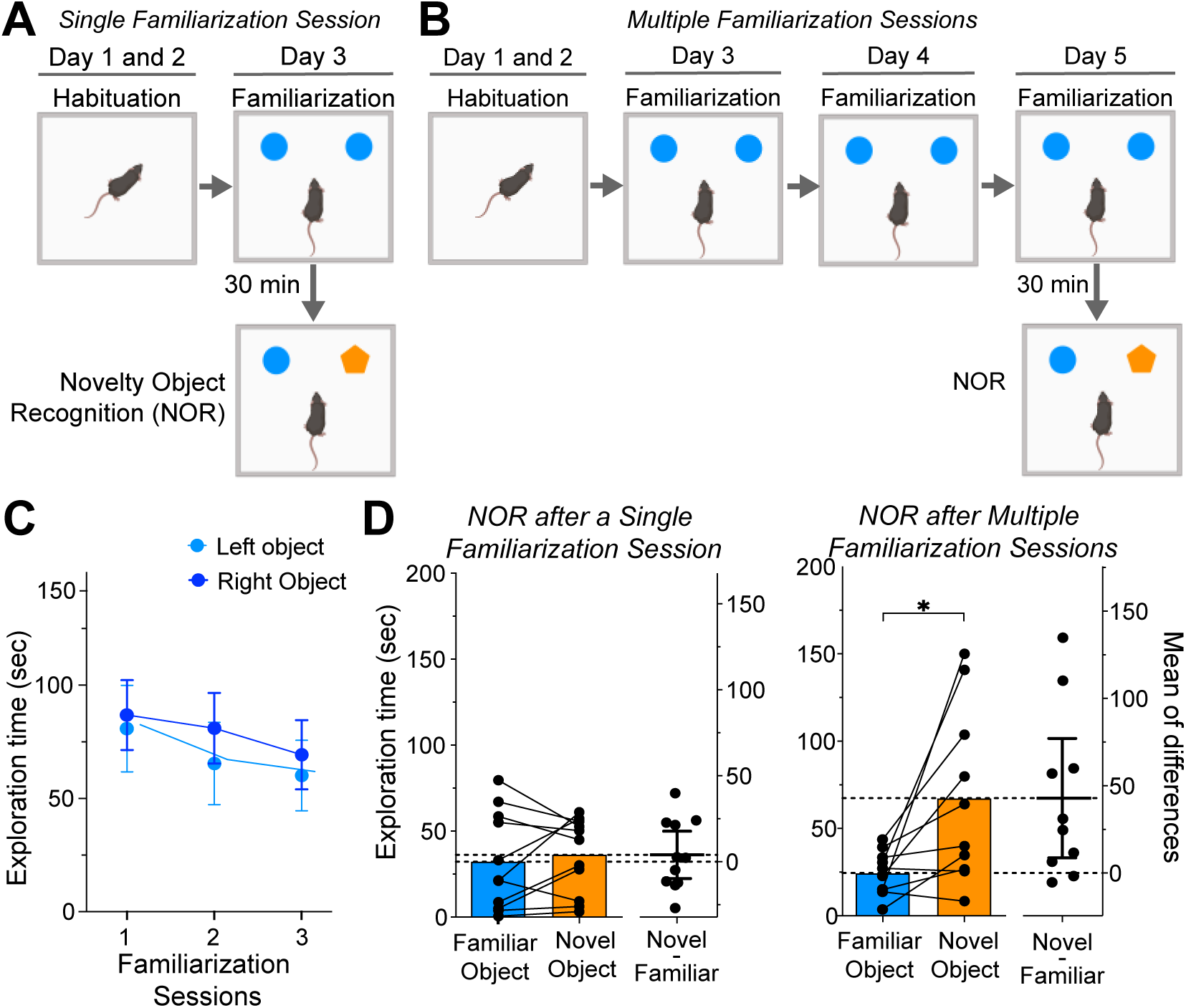
Impact of Familiarization Session Number on Novelty Object Recognition. (a) Single familiarization session NOR protocol schematic. (b) Multiple familiarization sessions NOR protocol schematic. (c) Graph of exploration time for two identical objects over three sessions, showing a minor decline (ANOVA, main effect of sessions, *P* = 0.0471). (d) Time exploring familiar and novel objects during NOR for groups exposed to single (left graph, n=11) and multiple (right graph, n=10) familiarization sessions; significant difference in exploration in multiple familiarization group (\**P* = 0.0098). The graphs’ right side represents the estimation plots showing mean differences (dots) and respective 95% confidence intervals.

The performance during the NOR session varied based on the number of familiarization sessions. Mice that underwent a single session displayed no preference between familiar and novel objects (**Figure 1D**, left; Wilcoxon matched pairs test; *P*=0.700). Conversely, mice subjected to multiple familiarization sessions showed a significant bias, spending more time with the novel object (**Figure 1D**, right; *P* = 0.009). Our data indicate that increasing familiarization sessions enhances novelty discrimination in mice.

### Multiple Familiarization Sessions are Associated with Reduced Dopamine Neuron Activity as Measured by c-Fos Expression

We investigated the extent of dopamine neuron activation triggered by single or multiple familiarization sessions in heterozygous TH-Flpo mice. This genotype was selected because it was used in the subsequent chemogenetic experiments. To assess changes in neuronal activity, we measured the expression of c-Fos, an accepted marker of neuronal activity (Yap and Greenberg, 2018). Mouse brains were harvested 90 minutes after single or multiple familiarization sessions (**Figure 2A**), coinciding with the peak c-Fos protein expression following object exploration (Tanimizu et al., 2018). We next quantified the number of TH+ neurons expressing c-Fos in the VTA in 3 brain sections located at −3.27, −3.51, and −3.63 mm from bregma (n=9 sections from 3 mice per group; **Figure 2B**). The immunoreactivity to TH was used as a dopamine neuron marker. We observed decreased dopamine neuron activity after extended familiarization, evidenced by a significant reduction in the relative number of TH+c-Fos+ neurons between single and multiple familiarization sessions (unpaired t-test; *P* = 0.0015; **Figure 2C**).

**Figure 2.**
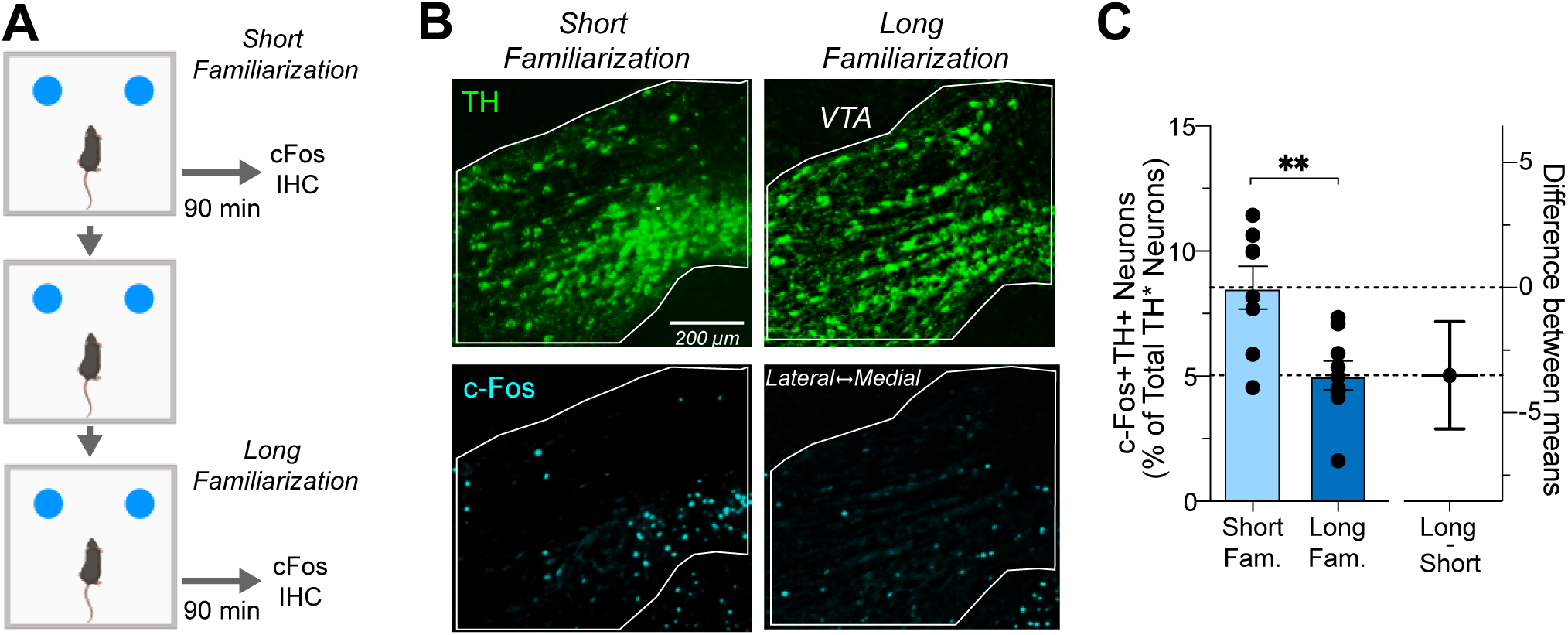
Dopamine Neuron c-Fos Expression Following Single or Multiple Familiarization Sessions. (a) Schedule for brain dissections post one or three familiarization sessions for c-Fos immunohistochemistry (IHC). (b) Photomicrographs of TH (upper panel) and c-Fos expression (lower panel) in the VTA after single or multiple sessions. (c) Left: Percentage of c-Fos+TH+ neurons relative to total TH+ neurons in VTA, showing a significant reduction post three sessions (n=9 brain sections in 3 mice per group, ** *P* = 0.0015). Right: Estimation plot showing effect size magnitude.

### Chemogenetic Inhibition of Dopamine Neurons During a Single Familiarization Session Improves Novelty Discrimination

To determine whether decreased dopamine neuron activity during training sessions contributes to familiarity, we examined if chemogenetic inhibition of these neurons during a single familiarization session could enhance novelty recognition, mimicking the effects observed in multiple familiarization sessions. Heterozygous TH-Flpo mice, expressing the inhibitory DREADD hM4Di selectively in dopamine neurons (**Figure 3A**), were divided into two groups: one receiving 1mg/kg CNO injections (n=17), and a control group given saline injections (n=13). In addition, mice that received mock surgeries and CNO injections were also tested to control for CNO effects on behavior (n=12). These groups underwent two rounds of NOR testing to evaluate the effects of inhibiting dopamine neuron activity during either the familiarization or NOR session (see **Figure 3B** schematic).

**Figure 3.**
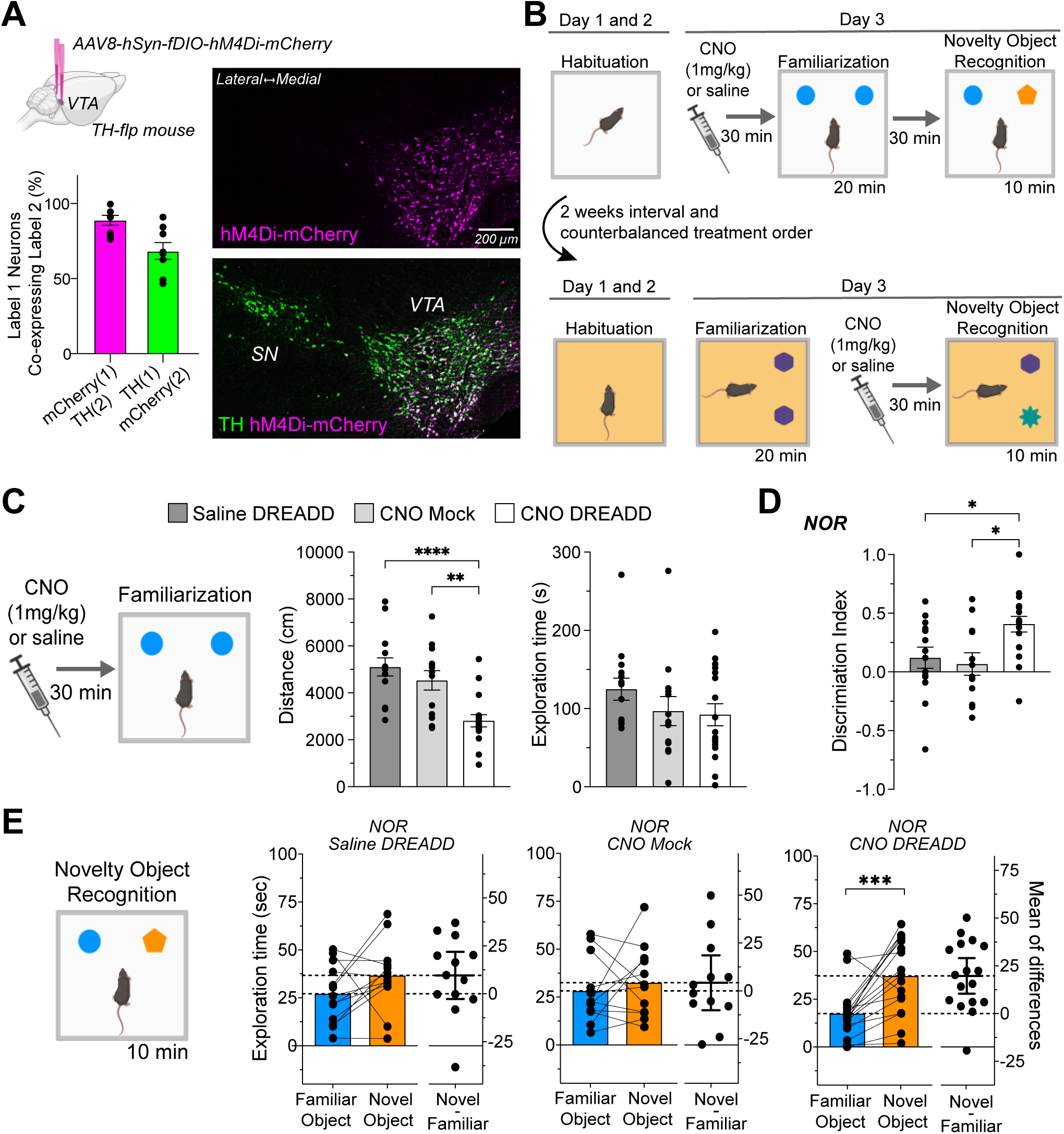
Effects of Chemogenetic Inhibition during Familiarization on Novel Object Recognition. (a) Brain diagram describing bilateral VTA DREADD virus injections. The graph below details the virus’s specificity (magenta bar: percentage of mCherry+ neurons co-expressing TH+) and its efficacy (green bar: percentage of TH+ neurons co-expressing mCherry+). Accompanying photomicrographs display DREADD virus (hM4Di-mCherry) expression in VTA TH+ neurons. (b) NOR paradigms schematic detailing CNO injection timing, either before or after the familiarization session, using a within-subject counterbalanced design. (c) Pre-familiarization CNO injections reduce travel distance (\*\*\*\**P* < 0.0001; **P = 0.032) without affecting object exploration time. (d) Discrimination index for object exploration during NOR. CNO DREADD group (n=17) showed increased discrimination compared to Saline DREADD (n=13, \**P* = 0.046) and CNO mock groups (n=12, \**P* = 0.018). (e) NOR session exploration times, with the CNO DREADD group displaying a pronounced novel object preference (\**P* = 0.0004).

Dopamine neuron inhibition during familiarization led to decreased locomotor activity (**Figure 3C**, left graph). A one-way ANOVA confirmed a treatment effect (F_(2, 41)_ = 12.70, P < 0.0001), and Bonferroni’s multiple comparison tests showed that the DREADD CNO group significantly differed from both control groups. Interestingly, this inhibition did not affect the average time exploring the objects (**Figure 3C**, right graph; Kruskal-Wallis test, *P* = 0.182). We subsequently assessed the effects on NOR performance and found that the DREADD CNO group had a significantly higher discrimination index (**Figure 3D**). A one-way ANOVA showed a significant treatment effect (F_(2, 40)_ = 5.188; *P* = 0.001), and Bonferroni’s tests revealed a significant difference between DREADD CNO and control groups. Furthermore, only the DREADD CNO group exhibited a significantly higher exploration of the novel object (**Figure 3E**; paired t-test; *P* = 0.0004). These results suggest that inhibiting dopamine neurons during familiarization enhances novelty discrimination in the subsequent NOR session.

### Chemogenetic Inhibition of Dopamine Neurons During the NOR session Does not Impact Performance

Finally, we assessed the impact of inhibiting dopamine neurons during the NOR session. This experiment determines if our manipulation affects memory retrieval of a previously encountered object. Additionally, it examines if the improved performance produced by the DREADDs during familiarization results from lingering effects on the NOR session.

During the NOR test, the DREADD CNO exhibited reduced locomotor activity (**Figure 4A**, left graph). One-way ANOVA verified the treatment’s effect (F_(2, 42)_= 9.733; *P* = 0.0003), and Bonferroni’s multiple comparison tests indicated a difference between the DREADD CNO and control groups. Notably, this suppression did not impact the average object exploration time (**Figure 4A**, right graph; (One-way ANOVA, F_(2, 42)_ = 0.257, *P* = 0.257). Moreover, the chemogenetic inhibition did not change the discrimination index (**Figure 4B**, One-way ANOVA, F_(2, 41)_ = 0.935, *P* = 0.400) or the exploration time for both novel and familiar objects (**Figure 4C**; paired t-tests; DREADD saline, *P* = 0.834; Mock CNO, *P* = 0.0791; DREADD CNO, *P* = 0.0698). Across all treatment groups, the time spent exploring the novel object was similar to that of the familiar object. The results suggest that inhibiting the activity of the dopamine neurons during NOR does not enhance memory recall of the previously encountered object and cannot account for the outcomes observed when inhibiting dopamine neurons during the familiarization session.

**Figure 4.**
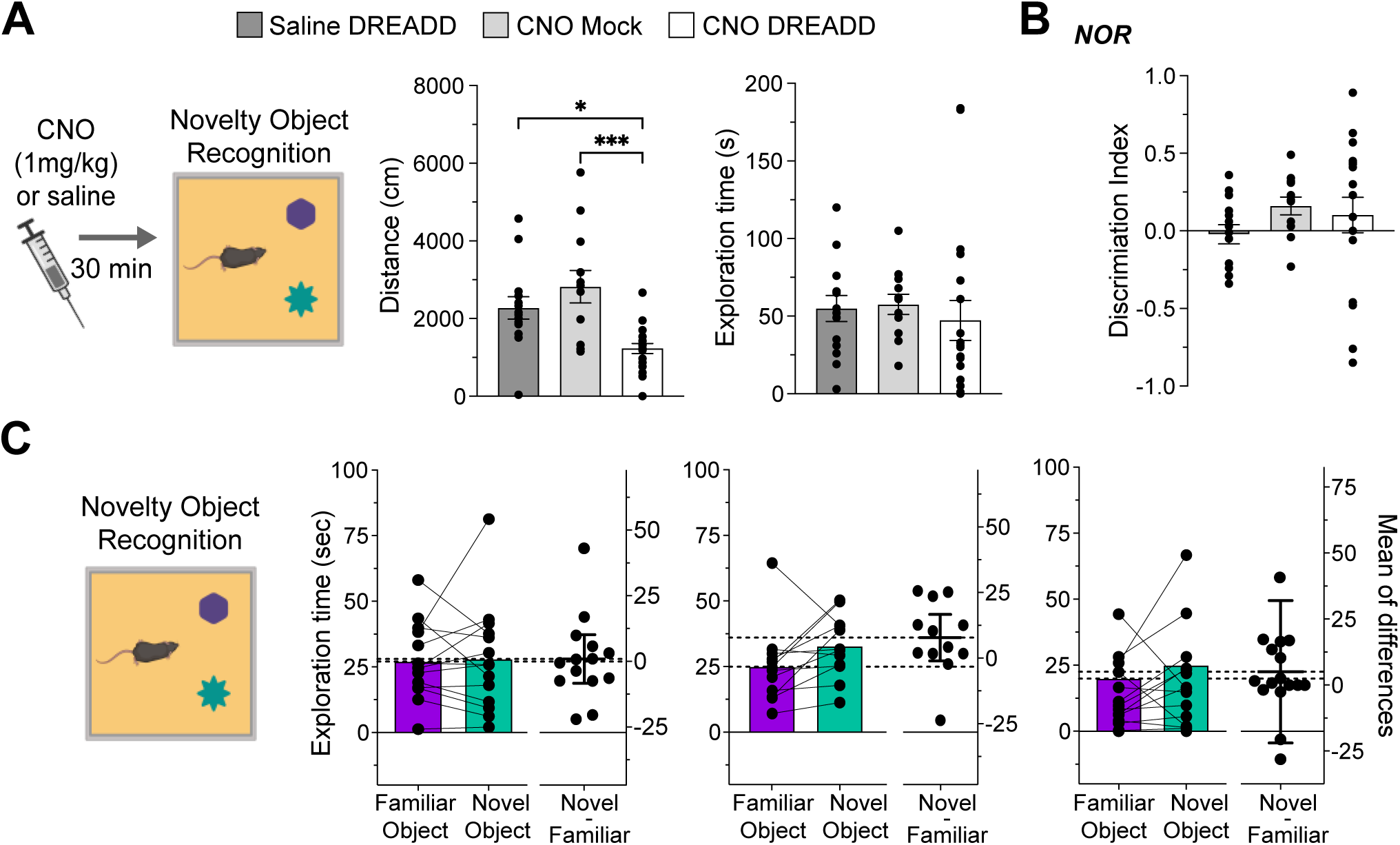
Effects of Chemogenetic Inhibition During NOR Session on Novelty Discrimination. (a) Pre-NOR session CNO injections reduce travel distance (*P = 0.017; ***P = 0.0004) but do not alter object exploration time (b) Discrimination index graph for the NOR session, with no significant difference between groups. (c) NOR session exploration times for both novel and familiar object; all groups exhibited similar exploration durations.

## Discussion

Our findings indicate that multiple sessions lead to decreased VTA dopamine neuron activation and enhanced subsequent novelty discrimination performance compared to a single familiarization session. Importantly, we presented evidence that the reduced activation of dopamine neurons during familiarization causally affects the subsequent discrimination between familiar and novel objects. By chemogenetically inhibiting these neurons during the exposure to a single familiarization session, we observed improved novelty recognition, effectively mimicking the effects of multiple familiarization sessions.

Why do multiple familiarization sessions produce robust novelty discrimination? Distinguishing between novel and familiar objects signifies successful memory consolidation and recall of the previously encountered object (Brown and Banks, 2015). Extended familiarization likely strengthens the memory consolidation of the familiar object (Albasser et al., 2009), and studies with rats have further corroborated that prolonged familiarization improves memory retention (Shimoda et al., 2021). Simultaneously, being exposed repeatedly to the same object without notable consequences leads mice to perceive the familiar object as both irrelevant and non-salient, thereby improving their discrimination of novel objects in subsequent encounters (Hornoiu et al., 2020; Osorio-Gómez et al., 2022). Therefore, multiple familiarization sessions could enhance novelty discrimination by strengthening memory consolidation and diminishing the perceived salience of the familiar object. Following this logic, the inability to distinguish between novel and familiar objects after a single familiarization might arise from either a weak recall of the object previously encountered or its sustained perception as salient. While our initial experiment does not discern between these two scenarios, our chemogenetic data suggest a role of salience in this process.

Chemogenetic inhibition of VTA dopamine neurons during either the familiarization or NOR session reduced locomotor activity, consistent with findings from our collaborators using the same virus (Runegaard et al., 2018). This finding not only underscores the established role of dopamine neurons in promoting locomotor activity (Iversen, 2016) but also validates the inhibitory effect of the DREADD virus in the current experiments. Dopamine neuron inhibition did not affect the time spent exploring objects during familiarization and NOR sessions, replicating previous findings (Gunaydin et al., 2014; Bariselli et al., 2018). Instead, dopamine shapes future responses to previously encountered objects, increasing novelty discrimination. The fact that improved performance is observed when inhibiting the neurons during familiarization and not during the NOR testing session indicates that the dopaminergic manipulation may impact memory acquisition rather than memory retrieval. However, neuropharmacology studies have shown that D1 and D2 receptor antagonism during familiarization does not affect memory acquisition (Besheer et al., 1999; Nagai et al., 2007). Furthermore, considering the known role of dopamine neurons in facilitating learning and memory (Steinberg et al., 2013; Nasser et al., 2017; Duszkiewicz et al., 2019), one might expect their inhibition to impair rather than enhance recognition memory.

Alternatively, the chemogenetic inhibition may impact the perceived salience of the familiar object in future interactions. There is compelling evidence that increased dopamine assigns salience to environmental stimuli (Berridge, 2007; Flagel et al., 2011; Kutlu et al., 2021). Our research contributes to this narrative by suggesting that the absence of VTA dopamine neuron activity enhances the perception of stimuli as irrelevant and non-salient. This conclusion aligns with studies using a latent inhibition paradigm, which demonstrated that inhibiting dopamine neurons decreases the perceived salience of cues and reduces their likelihood of being associated with either appetitive or aversive stimuli (Morrens et al., 2020; Kutlu et al., 2022). Finally, our hypothesis introduces a fresh perspective on why a single familiarization might yield weak novelty discrimination. Recognition memory forms during a single session, yet the previously encountered object retains its perceived salience during the NOR session, thereby increasing exploration. The inhibition of dopamine neurons diminishes this perceived salience, unmasking the underlying recognition memory. Thus, our results indicate that dopamine neurons play an essential role during familiarization by modulating the salience of memory content.

## Acknowledgements

This work was funded by a National Institute of Mental Health, R21 MH123926 awarded to SM, and the Lundbeck Foundation R231-2016-2481 awarded to ATS, and R276-2018-792 and R223-2016-261 awarded to UG.

## Author Contributions

Conceptualization: SM and SF; data curation: SM, SF, RK; formal analysis: SM, SF, RK, QH, JE, and JT; funding acquisition: SM; investigation: SM, SF, RK, QA, JE; methodology: SM, SF, RK, ATS, UG; project administration: SM, SF, RK; resources: SM, ATS, UG; software: SM, SF, RK, QA; supervision: SM, SF, RK, JT; validation: SM, SF, RK, QA; visualization: SM, SF, RK; writing: SM; and input from all authors.

## Data Accessibility

Data are available from the corresponding author upon request.

## Abbreviations

CNO: Clozapine-N-Oxide
DREADD: Designer Receptors Activated Only by Designer Drugs
IHC: Immunohistochemistry
NOR: Novel object recognition
VTA: Ventral Tegmental Area

